# Microfluidic immobilization and subcellular imaging of developing *Caenorhabditis elegans*

**DOI:** 10.1101/139915

**Authors:** Jordan Shivers, Sravanti Uppaluri, Clifford P. Brangwynne

**Author notes:** Present address: Department of Chemical and Biomolecular Engineering, Rice University, Houston, TX 77005, USA. Present address: School of Liberal Studies, Azim Premji University, Bangalore 560100, India (Sravanti Uppaluri). Author to whom correspondence should be addressed. Electronic mail.

## Abstract

*C. elegans* has been an essential model organism in the fields of developmental biology, neuroscience, and aging. However, these areas have been limited by our ability to visualize and track individual *C. elegans* worms, especially at the subcellular scale, over the course of their lifetime. Here we present a microfluidic device to culture individual *C. elegans* in parallel throughout post-embryonic development. The device allows for periodic mechanical immobilization of the worm, enabling 3D imaging at subcellular precision. The immobilization is sufficient to enable fluorescence recovery after photobleaching (FRAP) measurements on organelles and other substructures within the same specific cells, throughout larval development, without the use of chemical anesthetics. Using this device, we measure FRAP recovery of two nucleolar proteins in specific intestinal cells within the same worms during larval development. We show that these exhibit different fluorescence recovery as they grow, suggesting differential protein interactions during development. We anticipate that this device will help expand the possible uses of *C. elegans* as a model organism, enabling its use in addressing fundamental questions at the subcellular scale, including the role of phase transitions in driving spatiotemporal intracellular organization within multicellular organisms.

## INTRODUCTION

The nematode *Caenorhabditis elegans* is a powerful model organism due to its transparent body, simple nervous system, short life cycle and stereotyped development. These characteristics have enabled groundbreaking studies of physiological processes including apoptosis (Sulston and Horvitz 1977; Sulston et al. 1983), aging (Kenyon et al. 1993), the discovery of RNA interference (RNAi) (Tabara et al. 1998; Timmons and Fire 1998; Timmons et al. 2001), and studies of the formation and organization of intracellular structures (Lee et al. 2013; Weber and Brangwynne 2015; Feric et al. 2016). Significant progress has been made using developing *C. elegans* embryos to elucidate various aspects of intracellular structures, such as P granules and nucleoli (Brangwynne et al. 2009; Wang et al. 2014; Weber and Brangwynne 2015). However, much less is known about the assembly and function of such structures in post-embryonic development, due to the highly mobile nature of the developing worm larvae.

Following embryogenesis, post-hatch, growing worms become difficult to image at a subcellular level without a mechanism for immobilization. Chemical anesthetics such as levamisole hydrochloride are widely used, but given the lack of characterization of the effects of levamisole on development, and the potential impact on the biophysical properties of subcellular structures, an alternative method is needed. Other typical immobilization mechanisms include CO_2_ exposure (Bodri 2011) in a low-oxygen environment, as well as temperature reduction (Podbilewicz and Gruenbaum 2006), but the effects of these techniques on worm metabolism and specifically on post-embryonic development have not been well characterized. The specific mechanism underlying the anesthetic effect of CO_2_ on invertebrates is still poorly understood, though it has been proposed that it alters cell membrane permeability due to lowered intracellular pH levels (Nicolas and Sillans 1989; Chokshi et al. 2009). Previous experiments involving immobilizing worms via temperature reduction involve cooling worms to 4°C, and so achieving repeated immobilization during post-embryonic growth would require repeatedly cooling and warming the developing larvae (Chung et al. 2008). This is undesirable, as the effects of cyclic temperature variation on *C. elegans* development have not been characterized, while exposure to low temperature during larval stages has been shown to shorten worm lifespan (Zhang et al. 2015).

Mechanical immobilization within a microfluidic device offers a reversible, non-invasive option that is well suited to the flexible, cylindrical shape of *C. elegans*. Prior work has shown that stable and reversible immobilization can be achieved by pressurizing a flexible membrane (Mondal et al. 2012). This mechanism can be used to immobilize worms with stability comparable to that achieved with anesthetic methods, while maintaining normal growth and lifespan after immobilization (Gilleland et al. 2010). However, previous devices were designed to immobilize single worms of a specific size (L4 or Adult). Other microfluidic immobilization methods for physiologically active worms have been demonstrated (Hulme et al. 2007; Guo et al. 2008; Kim et al. 2013), but only one study has addressed the problem of periodically immobilizing and imaging individual worms on-chip during the entire duration of post-embryonic growth (Keil et al. 2016), and none have enabled automated loading of worms at the embryo stage. All previous devices require manually inserting individual, already-hatched worms. Inserting worms at the embryo stage is desirable because it could allow for imaging worms immediately after hatching, and loading liquid-suspended embryos would avoid the experimental difficulty of loading individual worms manually. Repeated stable immobilization of individual worms throughout their full lifetime is desirable to enable detailed studies of temporal changes in subcellular components within the same individuals while simultaneously gathering data from the full on-chip population.

We have previously demonstrated on-chip monitoring of growth over the entire course of *C. elegans* larval development, by utilizing fluid resistance to isolate embryos in individual chambers and constant flow-driven delivery of nutrients and waste removal (Brangwynne and Uppaluri 2015). However, without an immobilization mechanism, it is impossible to utilize techniques like 3D confocal imaging, fluorescence correlation spectroscopy (FCS), or FRAP, which require the worm to be completely immobile. Such techniques are crucial for the study of temporal changes in subcellular components. Indeed, quantitative confocal imaging and its variants have been essential to our understanding of organelle structure, size and formation (Brangwynne 2013; Weber and Brangwynne 2015; Berry et al. 2015; Feric et al. 2016; Uppaluri et al. 2016).

Over the last few years, there has been explosive interest in non-membrane bound organelles, due to the role they play in various physiological functions - from stress responses to ribosomal biogenesis (Brangwynne 2013; Hyman et al. 2014). These organelles are typically RNA and protein rich, and are referred to as ribonucleoprotein (RNP) bodies, and include stress granules, P granules, Cajal bodies, the nucleolus, among others (Brangwynne et al. 2015). Recent work suggests that these structures assemble by intracellular phase transitions, resulting in liquid droplets or gel-like assemblies (Lee et al. 2013; Zhu and Brangwynne 2015; Berry et al. 2015). The nucleolus exhibits a layered architecture, comprised of separate liquid phase compartments assembled around actively transcribing RNA polymerase 1. These include the dense fibrillar component (DFC), enriched in fibrillarin (FIB1), and the granular component which, in most organisms, is enriched in the abundant protein nucleophosmin (NPM1). Phase separation of these compartments is driven by differences in the miscibility and surface tension of the proteins comprising each compartment (Feric et al. 2016). Despite a number of studies using purified components in vitro, as well as cell culture models, few studies probe the phase behavior of RNP bodies within living multicellular organisms (Feric et al. 2016).

Here we describe a microfluidic device to culture and image physiologically active, isolated *C. elegans* individuals throughout all stages of development. The device is loaded with embryos, which are isolated into separate chambers and continuously supplied with a bacterial food source enabling complete growth and development. Worms can be immobilized by transient mechanical pressure by compression under a flexible PDMS membrane. Importantly the device architecture includes variable heights within the growth chamber allowing for imaging of worms of dramatically different sizes, from the first larval stage to adulthood. We show that the stability of this immobilization mechanism is consistent with that achieved with a typical anesthetic. We further demonstrate the ability to track changes in the morphology and biophysical properties of subcellular components in particular cells within the same worms, from the earliest larval stage to adulthood.

While preparing this manuscript, a method similar to ours was published by Keil et al. using the same membrane-pressurization mechanism of immobilization (Keil et al. 2016). Our device nonetheless offers a novel alternative for achieving immobilization during post-embryonic growth, with several distinctions. First, our device enables loading worms into the chambers as a single stream of embryos, which are automatically separated into the 8 chambers. This uniquely allows for imaging the worms immediately after hatching, or even during the embryo stage. In addition, our device offers the advantage of being comparatively simple to set up, as it requires only a single inlet and outlet tube and avoids the necessity of setting up an automated pressurization system to control each chamber, as pressurization is done by hand using a water-filled syringe with a valve and pressure gauge. Further, worms are immobilized linearly in our device, thus avoiding the complex image registration method required to straighten the worm images in the other device. Finally, we provide a quantitative measure of the immobilization stability achieved in our device, and we demonstrate a novel use of this immobilization method by performing FRAP on individual nucleolar components during development.

## METHODS AND MATERIALS

### Culture of C. elegans

*C. elegans* expressing intestinal FIB1::GFP and the *Xenopus laevis* protein NPM1::mCherry in all cells were grown at 20°C using standard methodology. For insertion into the device, plates of gravid adult worms were bleached using standard methods(Hope 1999) and embryos were washed repeatedly with M9 buffer and suspended in M9 for insertion into the device.

### Preparation of microfluidic devices

Devices were fabricated from polydimethylsiloxane (PDMS) using standard multilayer photolithography techniques (McDonald and Whitesides 2002). The flow layer consisted of 3 layers (10, 35, and 35 μm in height) and the compression layer consisted of a single 50 μm layer. These layers were fabricated separately and then bonded thermally. Prior to use, the compression layer (blue in Fig. 1a) was primed with water and the flow layer with M9 buffer. Embryos suspended in S-medium were then inserted into the device and trapped in individual chambers using the resistance-driven embryo trapping mechanism previously described by Uppaluri et al. (Brangwynne and Uppaluri 2015). Once each chamber contained a single embryo, S-medium was pumped into the device at 50 μL/hr for 12 hours, until all embryos hatched and became synchronized at the L1 growth stage. Then, S-medium supplemented with OP50 *E. coli* was flowed into the device at 50 μL/hr for the duration of the experiment, except for the brief immobilization and imaging steps, providing a continuous supply of food to the developing worms.

**FIG 1.**
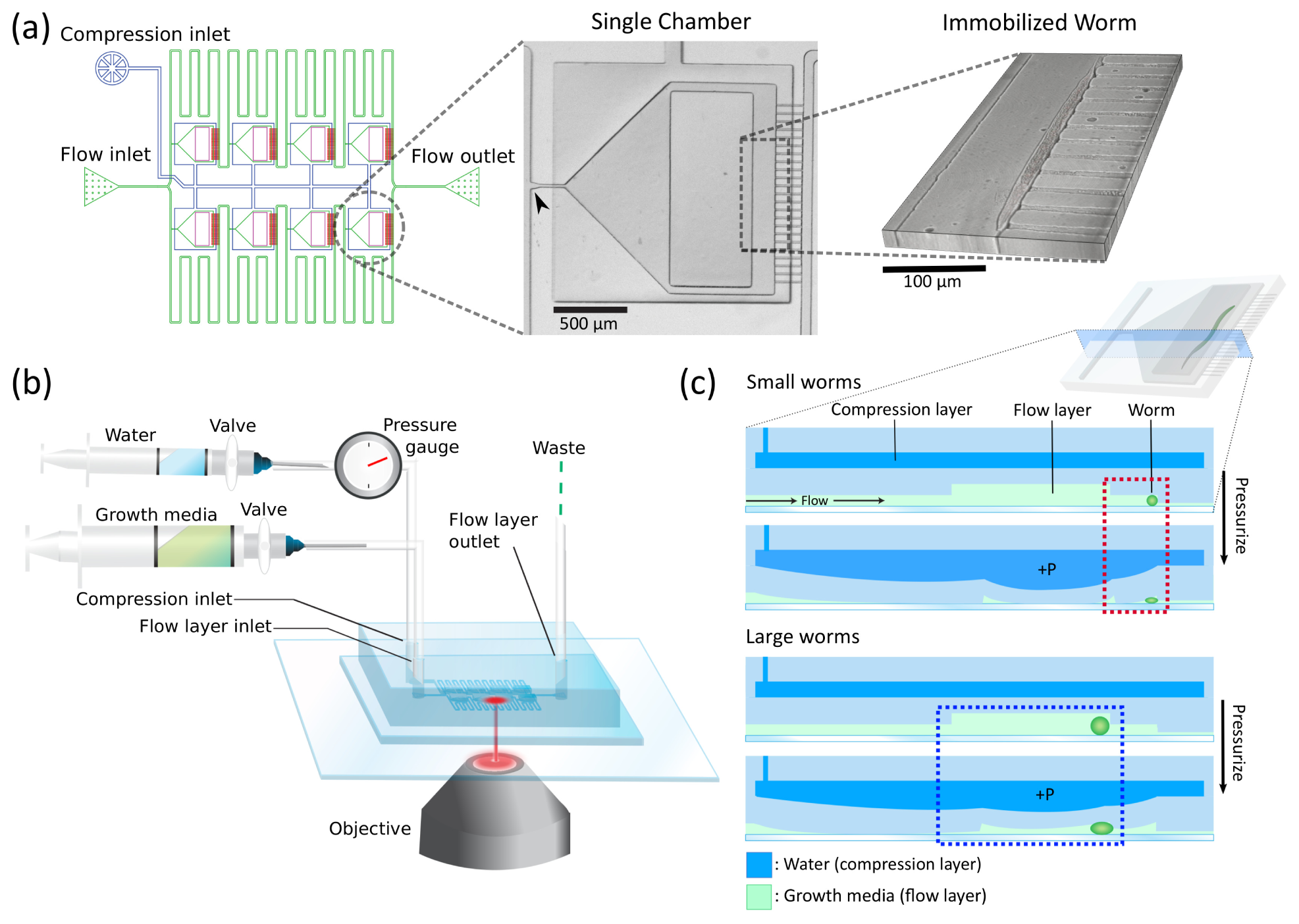
Design principles and immobilization mechanism. (a) Left: Schematic of 8-chambered device with flow layer (green) and compression layer (blue). Center: Top-down image of a single chamber within the fabricated device. Arrow designates embryo trap at the chamber entrance. Right: Z-stack of a young worm immobilized against outlet channels. (b) Schematic of experimental setup. (c) Cross section side-view of a single chamber illustrating the mechanism of compressive immobilization of smaller worms against the outlet channels (red dotted line) and larger worms in the central area (blue dotted line) with a thinner membrane.

### Immobilization and measurement of stability

The mechanism for immobilization involves pressurizing the compression layer (blue in Fig. 1a) to 15 psi using a water-filled syringe, as shown in Fig. 1b. This causes the membrane above each chamber to flex downward, as illustrated in Fig. 1c. The full membrane flexes down, and a thinner membrane in the center of the full membrane (blue dashed box in Fig. 1c) flexes further. Prior to pressurization, the flow rate in the device was briefly increased to force the worms to orient linearly, either against the outlet channels (small worms) or against the corner of the thin membrane (large worms). Once a pressure of 15 psi is achieved, a 2-way valve attached to the syringe is closed to maintain constant pressure. By design, worms can be immobilized in two areas: the 35 μm-height section against the outlet channels (red dotted line in Fig. 1c) for smaller worms, and a 70 μm-high section in the center, under the thinner membrane, for later larval stages. In both areas, worms are immobilized in a linear orientation perpendicular to the direction of flow, allowing for easy identification of cells of interest. We found that 15 psi was suitable to stably immobilize worms at all growth stages. Small worms were typically immobilized against the outlet channels, but they could also be immobilized in the center (as shown in the first image in Fig. 2a).

**FIG 2.**
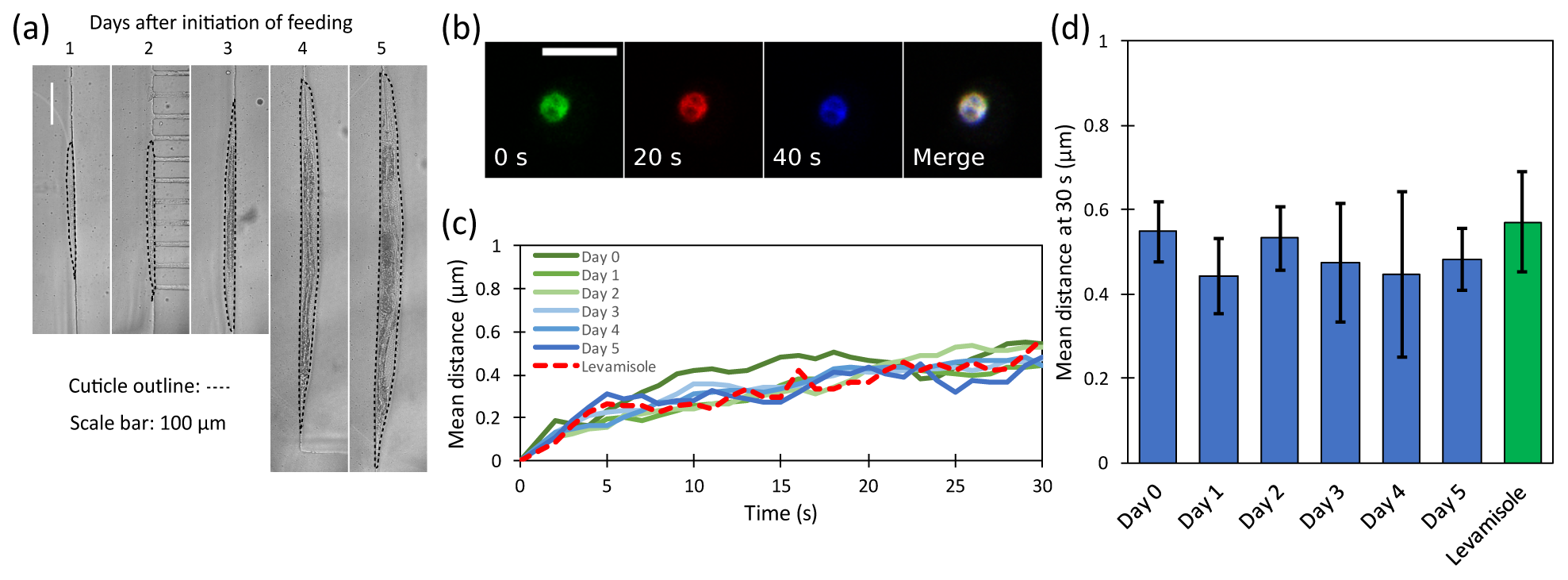
Stable immobilization at all growth stages. (a) Images of the same worm immobilized in a single chamber over 5 days. Images from days 1-2 show that small worms can be immobilized in the center as well as against the outlet channels. (b) Movement of a single nucleolus in an immobilized worm over 40 seconds, colored green (earliest), red, and blue (latest). The overlapping area appears white in the merged image, showing slight displacement over the recorded time period. Scale bar, 5 μm. (c) Average movement from the origin of several tracked nucleoli in 7 immobilized worms each day over a 5-day experiment is comparable to results for worms immobilized with levamisole. (d) Mean distance of nucleoli from original location after 30 seconds in 7 immobilized worms over 5-day period is comparable to results for worms immobilized with levamisole. Error bars represent S.E.M.

To measure immobilization stability, we recorded 30-second movies of several nucleoli in each immobilized worm each day and used particle tracking code (Crocker and Grier 1996; Blair and Dufresne 2008) to calculate the mean distance of all nucleoli from their original positions over time. For comparison, we recorded several movies of nucleoli in adult worms immobilized using levamisole and conducted the same analysis.

### Thrashing assay

Two devices were loaded with embryos and synchronized. For each chamber in each device, 1-minute movies of worms were recorded at 10 fps and side-to-side head motions were counted every 24 hours according to the body-bending or “thrashing” assay (Nawa and Matsuoka 2012). Worms in the control device were never immobilized, whereas worms in the “compression” device were subjected to an hour of immobilization at 15 psi per day. Worms were allowed 24 hours to recover from immobilization, and each thrashing measurement was taken immediately before the next immobilization cycle.

### Measurement of fluorescence recovery of nucleolar proteins

FRAP experiments were conducted using a Nikon A1 laser scanning confocal microscope equipped with a 60x oil immersion objective. Fluorescence recovery of a bleached region of interest (ROI) of 1-μm radius in nucleoli in the second intestinal ring (int-2) was recorded. The second intestinal ring was chosen because it contains two cells with large nucleoli that do not divide after initial cell differentiation. Fluorescence data is double normalized to correct for photobleaching and total fluorescence loss in the bleached nucleolus(Phair et al. 2003). This involves measuring average fluorescence in 3 regions as a function of time, t: a bleach ROI (the area in the nucleolus to which a bleaching pulse is applied) with intensity *I*(*t*), an ROI encompassing the whole nucleolus with average intensity *T*(*t*), and a background ROI with intensity *BG*(*t*). Average pre-bleach values *I*_*pre*_, *T_pre_*, and *BG_pre_* are calculated. The normalized fluorescence intensity *I_norm_*(*t*) is then calculated as follows:

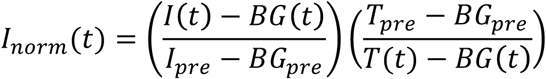

With this normalization method, full recovery is expected unless an immobile fraction is present. More discussion of the normalization method is available in Supplementary Information. Normalized fluorescence recovery measurements were fit to an exponential function of the form *f*(*t*) = *A*(1 − *e*^−*t*/*τ*^) from which we determined the recovery timescale *τ*.

## RESULTS AND DISCUSSION

With this device, embryos are trapped by hydrodynamic resistance in isolated chambers (8 in total) as in Uppaluri et al. (Brangwynne and Uppaluri 2015), allowed to hatch and synchronize to the L1 stage, and then periodically immobilized within these chambers throughout larval development. The design utilizes a membrane of varied thickness, with separate sections for immobilization during earlier and later larval stages, accounting for the significant changes in body size (~10-fold increase in worm diameter) of growing worms. The size and thickness of these sections was optimized to allow for stable imaging of worms of all sizes, allowing for 3D confocal imaging and FRAP with high spatio-temporal resolution at all larval stages. In the following sections, we characterize the functionality of the device and demonstrate its capability.

### Stable immobilization at all larval stages

Images of a single worm immobilized in the device periodically from L1 stage to adulthood are shown in Fig 2a. The device enables stable linearly-oriented immobilization of worms at all larval stages. Young worms (L1/L2-stage) are typically immobilized against the outlet channels, but they can also be immobilized in the center, as shown in the leftmost image in Fig 2a. At all larval stages, immobilization stability was sufficient for imaging subcellular structures, capturing Z-stacks to reconstruct 3D images, and performing FRAP experiments on individual nucleoli.

Worms immobilized within the device at all developmental stages exhibit a very high immobilization stability, with a mean displacement of significantly less than 1 μm, even after >30 seconds of sustained immobilization; this is comparable to the immobilization determined for worms immobilized with the anesthetic levamisole, as shown in Fig 2c-d. We further verify that the mechanical immobilization within the device does not impact the health of the worms, using a thrashing assay (Nawa and Matsuoka 2012) to measure body movements (Fig. 3a). As shown in Fig. 3b, worms cultured in the devices with and without immobilization do not exhibit any differences in body movements. This is consistent with prior mechanical immobilization results which determined that such immobilization did not appear to affect worm lifespan(Gilleland et al. 2010; Keil et al. 2016).

**FIG 3.**
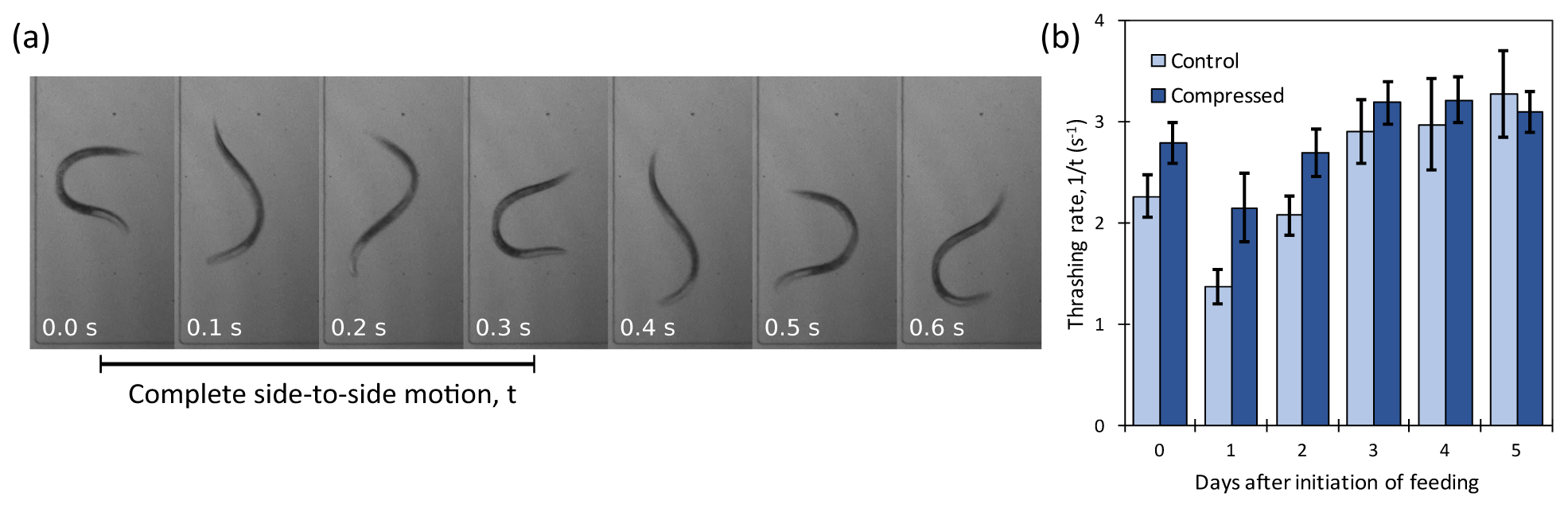
Immobilization does not affect worm motility. (a) Time lapse images showing thrashing of a freely swimming worm in a single chamber in the device. Two complete side-to-side motions are visible over 0.6 seconds. (b) Influence of 1 hour of immobilization in the device per day on the side-to-side thrashing rate of worms over 5 days, compared to worms in a control device not subjected to compression. Immobilization does not appear to affect worm motility. Error bars represent S.E.M.

### Stable imaging of subcellular structures in developing larvae

We observe morphological changes in the architecture of specific intestinal nucleoli in the same developing worms, as shown in Fig. 4b. *C. elegans* does not contain a homologue of the nucleolar protein nucleophosmin (NPM1), but exogenous GFP tagged NPM1 localizes to an outer GC layer, giving rise to a “core-shell” configuration, as in other organisms. However, older worms exhibited more complex configurations and morphologies with one or more inner cores of NPM1, surrounded by a layer of FIB1, which was surrounded by another layer of NPM1. These configurations are reminiscent of multiple-emulsion droplets produced using microfluidic techniques(Abate et al. 2011), and support the idea that the FIB1 and NPM1 components of nucleoli are coexisting liquid phases separating due to varied surface tension and miscibility(Feric et al. 2016).

**FIG. 4.**
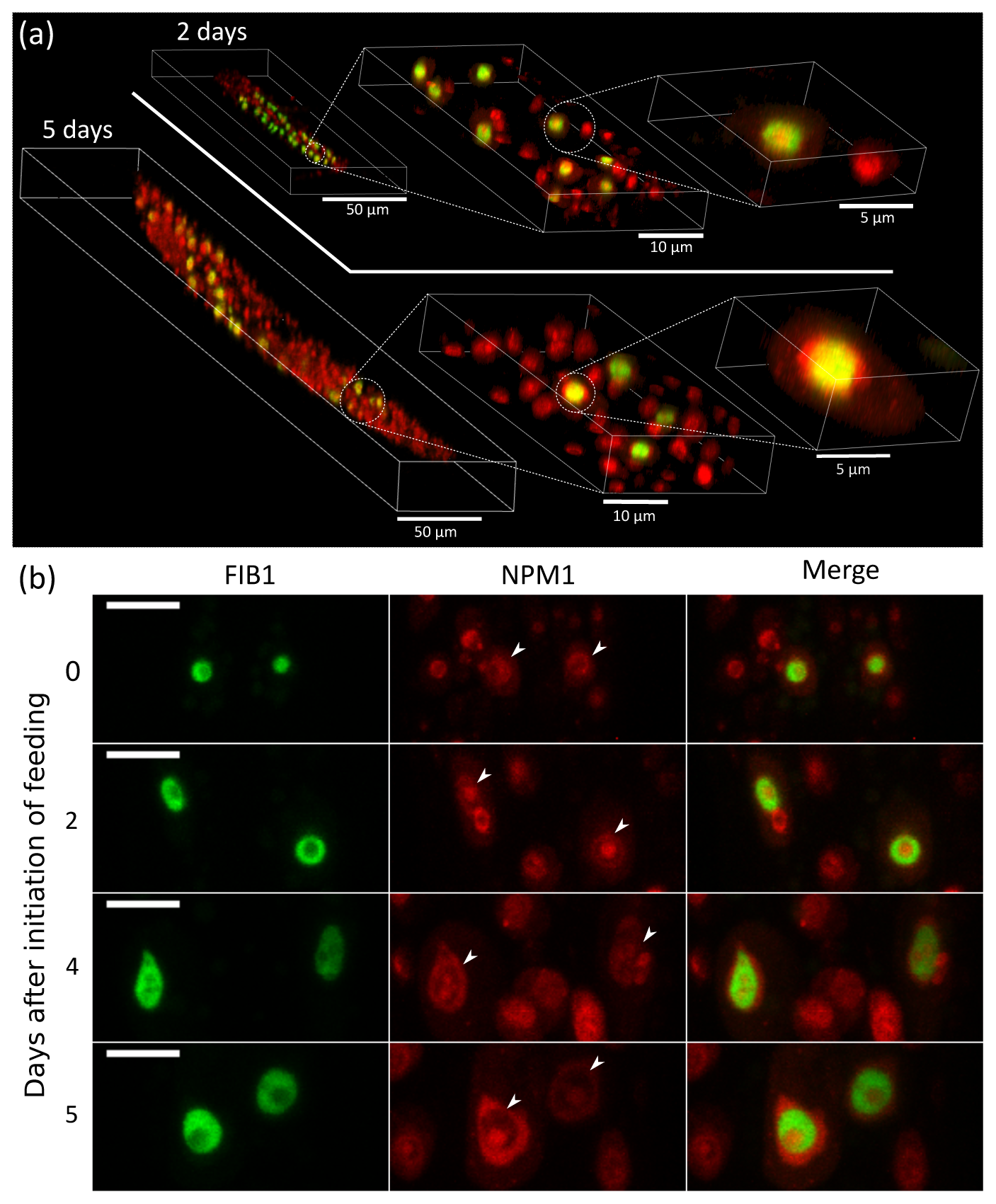
On-chip high-resolution imaging of the same cells in the same worms throughout larval development. (a) 3D renderings of Z-stacks of the same worm at 2 days (above white line) and 5 days after initiation of feeding. Left: Full body. Middle: Cells in the vicinity of the int-2 cell ring. Right: Single int-2 nucleolus. Compressive immobilization in the device allows for detailed imaging and reconstruction of cellular structure throughout larval development. (b) Maximum intensity projections of Z-stacks of the same int-2 cell nucleoli in a single worm in the device during growth. Changes in the size and internal organization of the nucleoli are observed. Arrows in the middle column point to int-2 nucleoli. Scale bar, 5 μm.

The immobilization stability achieved with this device enables capturing *Z*-stacks of the whole worm or specific cells in the same physiologically active worm throughout its growth. Figure 4a shows 3D reconstruction of a particular worm and its int-2 nucleoli at 2 days (L2/L3 stage) and 5 days (adult) of growth, demonstrating the ability to resolve structural changes at the whole-organism and subcellular level throughout larval growth using the device. Because worms are isolated in separate chambers, such structural changes can be simultaneously observed for many distinct worms in the device over their lifetime.

### Measuring temporal changes in the dynamics of nucleoli in developing worms

Using the device, we observed 7 isolated individual worms over 5 days post-hatching synchronization, until egg laying and performed FRAP on the same int-2 nucleoli. We observed a distinct increase in the timescale of fluorescence recovery of the FIB1 component of intestinal nucleoli over this period (Fig. 5a). We also sought to examine proteins enriched in the outer granular component (GC) layer of nucleoli. In contrast to the behavior of FIB1, NPM1 showed no change in the FRAP recovery time at different developmental stages (Fig. 5b). These results are consistent with prior work which suggests that droplets of FIB1, but not NPM1, appear to age *in vivo* in liquid droplets and *in vivo* in *X. laevis* oocytes and *C. elegans* worms(Feric et al. 2016). Fitting the fluorescence recovery data to an exponential, we find that the recovery timescale *τ* increases roughly threefold for FIB1 and remains constant for NPM1 (Fig. 5d). All results showed complete recovery, which by the double normalization method indicates a negligible immobile fraction of FIB1 or NPM1 within the nucleolus (Phair et al. 2003). Our results support the idea that the FIB1-rich DFC undergoes time-dependent changes to its viscoelastic properties. Thus, we demonstrate that the device enables immobilization at all larval stages sufficient to repeatedly perform stable confocal microscopy and FRAP, and thus resolve time-dependent changes in the material properties of the same cells in the same worms during larval development.

**FIG 5.**
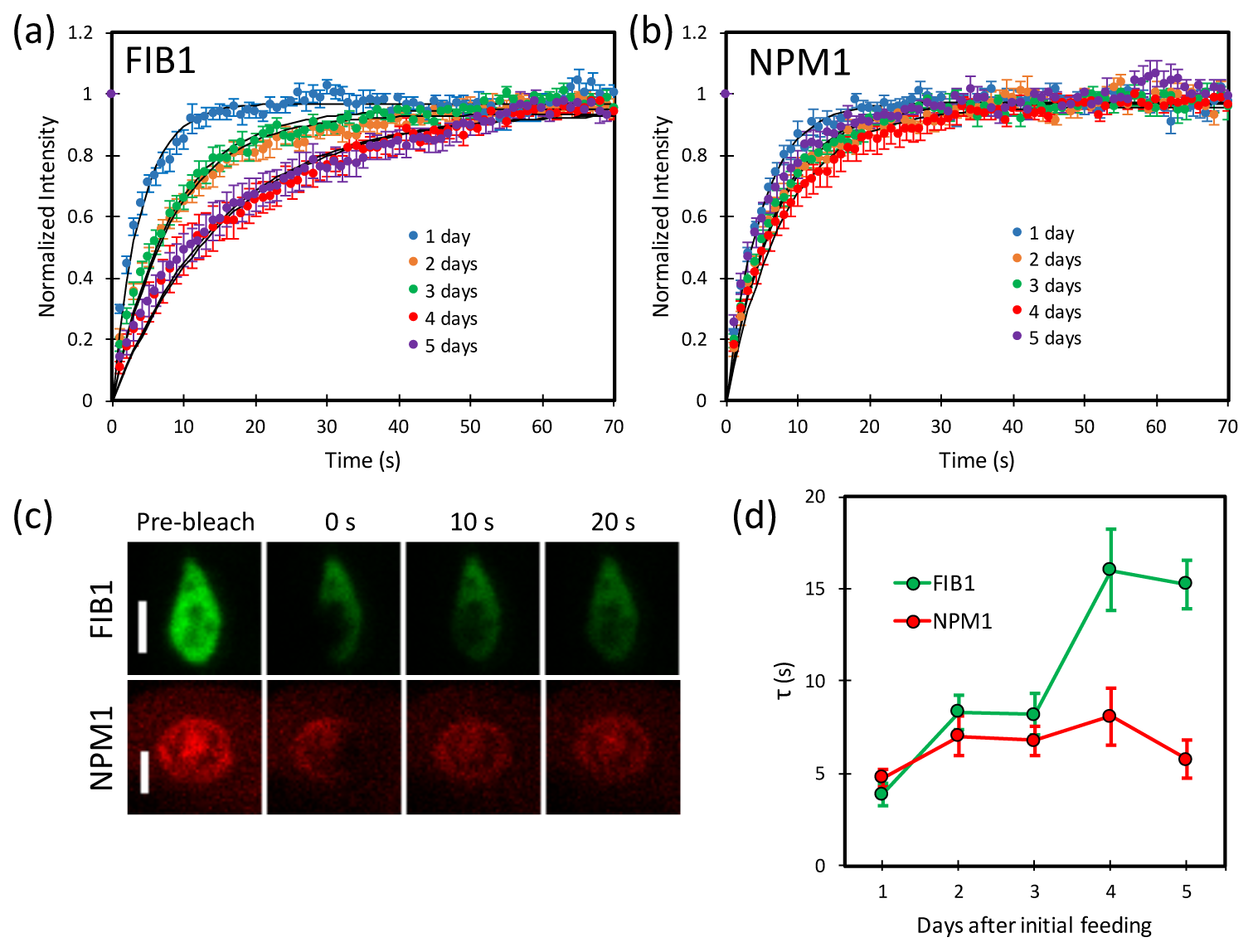
On-chip FRAP measurement of the same cells in the same worms throughout larval development. (a) Fluorescence recovery of FIB1 component and (b) NPM1 components in int-2 nucleoli in the same worms over 5 days show consistently fast recovery for NPM1 but a slowing in the rate of recovery of FIB1 as larval development progresses. Curves represent least-squares fit to *f*(*t*) = *A*(1 − exp(−t/*τ*)) and error bars represent standard error of the mean. (c) Representative time-lapses of FRAP in the FIB1 and NPM1 components of two nucleoli in a Day 4 worm. Rapid recovery of NPM1 is observed, compared to slower recovery of FIB1. Scale bars, 2 μm. (d) Calculated recovery timescale *τ* for FIB1 and NPM1 components over 5 days of larval development. Error bars represent S.E.M.

## CONCLUSIONS

We present a microfluidic device that allows for culture and stable, periodic immobilization of individual, trackable *C. elegans* worms from the earliest larval stage into adulthood. We demonstrate that the device achieves immobilization stability that is comparable to that achieved with the anesthetic levamisole, and we use the device to track morphological and biophysical changes in the components of specific intestinal nucleoli within the same worms throughout their growth. The device could be readily scaled up to include more chambers in series, which would enable even larger population-level studies. With stability sufficient to conduct 3D confocal imaging of developing worms at all growth stages, this microfluidic approach could potentially be applied to the study of any number of dynamic properties of developing C. elegans, as it enables the measurement of temporal changes at the subcellular level, in addition to 3D imaging of the entire organism. This device thus enables linking assembly and function at several levels of biological organization, from organelle to organism, in individual worms via quantitative imaging. As a unique but complementary approach to recent published work (Keil et al. 2016), our device offers a robust, technically simple and easy-to-implement alternative, with the added benefits of linear immobilization and the ability to automatically load individual embryos into isolated chambers.

## SUPPLEMENTARY MATERIAL

See supplementary material for detailed description of the FRAP normalization method and AutoCAD schematic for the device.

## ACKNOWLEDGEMENTS

We would like to thank Carlos Chen and the members of the C.P.B. lab for helpful discussions. We also thank Saurabh Vyawahare at the Princeton Microfluidics Facility. Some strains were provided by the CGC, which is funded by the NIH Office of Research Infrastructure Programs (P40 OD010440). This work was supported by the NIH Director’s New Innovator Award (1DP2GM105437-01) and the Searle Scholars Program (12-SSP-217). J.S. was supported in part by Princeton University’s Lidow Senior Thesis Fund.

